# Transcriptome-wide analysis of pseudouridylation in *Drosophila melanogaster*

**DOI:** 10.1101/2022.09.16.507327

**Authors:** Wan Song, Ram Podicheti, Douglas B. Rusch, W. Daniel Tracey

## Abstract

Pseudouridine (Psi) is one of the most frequent post-transcriptional modification of RNA. Enzymatic Psi modification occurs on rRNA, snRNA, snoRNA, tRNA, non-coding RNA and has recently been discovered on mRNA. Transcriptome-wide detection of Psi (Psi-seq) has yet to be performed for the widely studied model organism *Drosophila melanogaster*. Here, we optimized Psi-seq analysis for this species and have identified thousands of Psi modifications throughout the female fly head transcriptome. We find that Psi is widespread on both cellular and mitochondrial rRNAs. In addition, more than a thousand Psi sites were found on mRNAs. When pseudouridylated, mRNAs frequently had many Psi sites. Many mRNA Psi sites are present in genes encoding for ribosomal proteins, and many are found in mitochondrial encoded RNAs, further implicating the importance of pseudouridylation for ribosome and mitochondrial function. The 7SLRNA of the signal recognition particle is the non-coding RNA most enriched for Psi. The three mRNAs most enriched for Psi encode highly-expressed yolk proteins (Yp1, Yp2, Yp3). By comparing the pseudouridine profiles in the *RluA-2* mutant and the *w*^*1118*^ control genotype, we identified Psi sites that were missing in the mutant RNA as potential RluA-2 targets. Finally, differential gene expression analysis of the mutant transcriptome indicates a major impact of loss of RluA-2 on the ribosome and translational machinery.

## Introduction

Pseudouridylation is one of the earliest discovered and most abundant post-transcriptional RNA modifications. Pseudouridine (Psi), the C5-glycoside isomer of uridine occurs at specific nucleotide residues in rRNAs, tRNAs, snRNAs, snoRNAs and other ncRNAs (Ge and Yu 2013) and more recently Psi has also been found in mRNA (Carlile et al. 2014; Schwartz et al. 2014; Lovejoy et al. 2014; Li et al. 2015). Psi sites within RNAs are thought to have various functions depending on the molecule. Psi sites in tRNAs are present in the anticodon stem-loop, D-stem and other conserved sites and contribute to the stabilization of tertiary structure. In rRNA the known Psi sites are concentrated near the decoding site, the peptidyl transferase center, and at sites of interaction with ribosomal proteins, where they are thought to be important for ribosome assembly and protein synthesis (Charette and Gray 2000). In the non-coding RNA (ncRNA) of telomerase a functionally important Psi residue is located in a region involved in telomerase reverse transcriptase binding (Penzo et al. 2017). Incorporation of Psi into therapeutic mRNAs results in a reduced immune response to the RNA and enhanced protein expression (Kauffman et al. 2016).

Experimental detection of pseudouridine residues has been hindered and complicated by the fact that Psi and uridine share the same molecular mass and similar base-pairing properties upon reverse transcription. The most frequently used method to detect this nucleoside is to treat RNA with cyclohexyl-N’-(2-morpholinoethyl)-carboiimide metho-p-toluene sulfonate (CMCT). CMCT reacts with uridine-like and guanosine-like nucleotides and subsequent alkaline treatment of the reacted RNA hydrolyzes CMC adducts to G and U but does not hydrolyze CMC-Psi mono-adducts. The bulky CMC moiety blocks primer extension (Ho and Gilham 1967) and this block may be directly detected with polyacrylamide gel electrophoresis and autoradiography in highly abundant RNA species such as rRNA. Pioneering work by the Ofengand group used this reaction to detect highly conserved Psi sites in rRNA in species ranging from prokaryotes to eukaryotes (Bakin and Ofengand 1993).

More recently next generation RNA sequencing approaches have been adopted for detection of pseudouridines in lower abundance transcripts. The majority of these methods also depend on CMCT treatment since the CMC adduct conjugated to Psi prevents read-through by reverse transcriptase (RT). Bioinformatic analyses of CMC treated libraries identifies sites of pseudouridylation where reverse transcription is more frequently terminated relative to reads in untreated libraries. This Psi-seq approach (also known as “pseudo-seq”, “Ψ-seq”, or “CeU-Seq”) has been employed for global transcriptome-wide identification of pseudouridylation sites in yeast (Carlilie et al. 2014, Schwartz et al. 2014, Lovejoy et al. 2014), human cell lines (Schwartz et al. 2014, Carlilie et al. 2014, Li et al. 2015), mouse (Li et al. 2015), *Toxoplasma gondii* (Nakamoto et al. 2017) and *Arabidopsis thaliana* (Sun et al. 2019).

The post-transcriptional isomerization of uridine to pseudouridine is catalyzed by six families of pseudouridine synthases (TruA, TruB, TruD, RsuA, RluA and Pus10). The archaeal and eukaryotic TruB homologues function with auxiliary guide RNAs while the rest act as stand-alone enzymes (Hamma and Ferre-D’Amare 2006; Ofengand 2002). Some pseudouridine synthases have highly conserved and specific substrates whereas others show more broad and complex targets (Rintala-Dempsey and Kothe 2017). In the *Drosophila melanogaster* genome, there are nine proteins with an annotated pseudouridine synthase domain. *Drosophila* Nucleolar protein 60B (Nop60B), also known as *minifly* (*mfl*), is a well-studied homolog of human box H/ACA RNP dyskerin (mouse NAP57 and yeast Cbf5). Nop60B/Mfl is essential for *Drosophila* viability and fertility as it is required for ribosome biogenesis and the *mfl* mutant has reduced pseudouridylation at several sites of 28S and 18S rRNA (Phillips et al. 1998; Giordano et al. 1999; Breznak et al. 2022). Two other pseudouridine synthases (RluA-1 and RluA-2) have been studied for their roles in the negative regulation of nociception (Song et al. 2020). RluA-1 is primarily expressed in multidendritic sensory neurons including the larval nociceptors whereas RluA-2 is expressed ubiquitously. RluA-1 and RluA-2 belong to RluA family which have complex substrate specificity in yeast and bacteria (Hoang et al. 2006).

Here, we have applied the Psi-seq technique to identify sites of pseudouridylation in *D. melanogaster* female head transcriptome. In addition, we have investigated the pseudouridine and transcriptional profile in animals lacking the ubiquitously expressed RluA family member RluA-2. Our results suggest a major role for RluA-2 for pseudouridylation across a variety of RNA classes as well as large changes in gene expression that occur in animals lacking RluA-2 function.

## Results

### Optimization of Psi-seq library preparation conditions for efficient detection of putative pseudouridine sites

We began our experiments with optimization of a previously used strategy (Schwartz et al. 2014) and applied the optimized technique to *Drosophila* head RNA. As an internal control to monitor the effectiveness of the treatment and efficiency of the Psi detection, we spiked in a 150bp *in vitro* transcribed RNA oligonucleotide containing a single pseudouridine at position 43 (Schwartz et al. 2014) at an early stage of library preparation. In pilot experiments we identified experimental treatment conditions that allowed for robust calling of the pseudouridine of the RNA spike-in (**Figure S1**, details in Materials and Methods). Our RNA samples were split into two equal halves immediately after polyA enrichment with one half subjected to CMCT treatment and the other to a mock treatment. This was followed by a ligation of an RNA adapter, an adapter-specific RT reaction, then a second ligation of a DNA adapter. The samples were then barcoded, amplified, and pooled for Illumina sequencing (**Figure 1A**).

**Figure 1.**
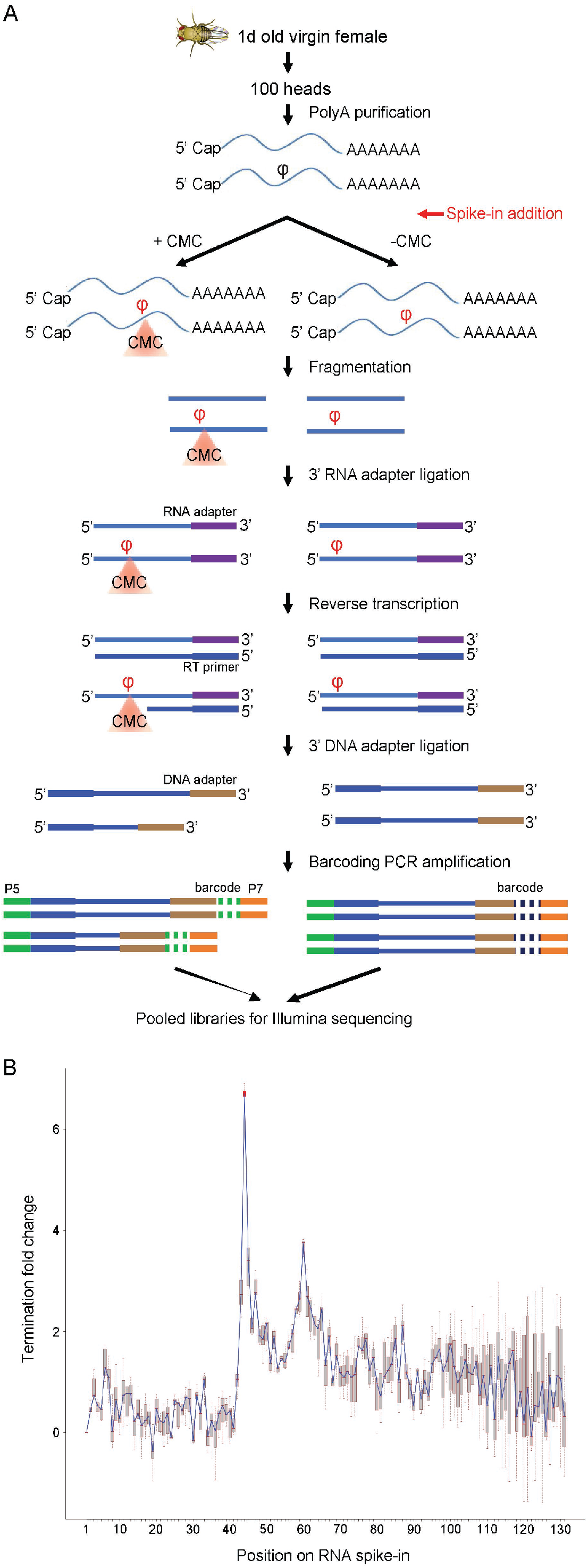
**A**. Schematic of *Drosophila melanogaster* Psi-seq library construction. RNA was extracted from heads of unmated female flies. An *in vitro* transcribed RNA spike-in containing a single pseudouridine at position 43 was added into the RNA samples immediately after polyA selection and the optimized CMCT derivatization step (**Figure S1**) was carried out prior to RNA fragmentation and subsequent library preparation. Libraries were barcoded and pooled for the final Illumina sequencing. **B**. A connected box plot showing the termination fold change (log2) on each of the nucleic acid position along the synthetic RNA spike-in added to the genetic background *w*^*1118*^ libraries (n=7) used for pseudouridine mapping. With the optimized CMCT treatment (**Figure S1**), the termination fold change peak at the only Psi (position 43) of the RNA spike-in (highlighted with the red box plot) stands out among all the other positions.

The RNA samples we used for the mapping of pseudouridylation were isolated from the heads of one day old virgin female flies of an isogenized *w*^*1118*^ (iso*w*^*1118*^) strain. This genetic background was previously used for behavioral characterization of *RluA-1* and *RluA-2* mutants (Song et al. 2020) and we focused on the head tissue since it would be enriched for genes expressed in the nervous system which are of greatest interest to us given the nociception phenotypes in the mutants. To assess repeatability, we carried out two sets of experiments. In experiment 1 (Exp 1) we prepared, sequenced, and analyzed three pairs of *w*^*1118*^ libraries (with each pair containing a CMCT and mock treatment). In experiment 2 (Exp 2) we performed the procedures on four pairs of *w*^*1118*^ and four pairs of *RluA-2*^*del-FRT*^ mutant (**Table S1 in File S1**). An average of 91% of the paired reads from the 22 libraries had both mates cleared through quality filters and more than 87% of the reads could be mapped concordantly (**Table S2 in File S1**), confirming the high quality of the sequencing reads for the libraries from both experiments.

To call potential Psi sites we determined a *termination ratio* for all nucleotide sites in the sequenced libraries. The termination ratio was computed as the ratio of read counts that terminated at a specific site in CMC treated samples and the total coverage in mock treated samples. Since the CMC treatment blocks reverse transcription adjacent to a Psi site, a higher termination ratio in the CMC library compared to the corresponding mock library is an indicator of the location of a potential Psi residue. We then used the log2 of termination ratio in the paired CMC and mock libraries to define *termination fold change*. Our bioinformatics pipeline, when applied to the synthetic spike-in RNA in our experiments resulted in termination fold change at the expected position adjacent to the Psi site that was greater than 6 across all pairs of CMC and mock libraries (**Table S2 in File S1**). A termination fold change greater than 6 represents an average absolute fold change greater than 64 (since log_2_(64) = 6) (**Figure 1B**). This result for the RNA spike-in indicates a highly efficient CMCT treatment which produced strong termination adjacent to the Psi site.

### Pseudouridylation of Drosophila ribosomal RNAs

Previous studies have mostly employed subjectively chosen thresholds for Psi site calling, i.e. a combination of “Psi ratio” (equivalent to our “termination ratio”) > 0.1 and “Psi fold change” (equivalent to our “termination fold change”) of >3 were used to identify the candidate pseudouridine sites in yeast and human cell lines (Schwartz et al. 2014). We found that applying the same thresholds to our data resulted in the calling of an extremely high number of sites. For instance, if we applied the thresholds of Schwartz et al (2014) we identified over 200,000 sites across seven *w*^*1118*^ libraries. In addition, we found that reproducibility of sites called using this thresholding method was low when comparing across replicates. The very high number of sites using the thresholding and the lack of reproducibility across samples caused us to question the robustness of this calling method for our experimental conditions.

To overcome this limitation, we analyzed the potential Psi sites according to the statistics of the termination ratios that we observed. To do this we applied a two-proportion z test to evaluate whether the termination ratio at a specific site in the CMC treated sample was significantly higher than in the mocked treated sample. Sites with an associated FDR<0.05 are statistically more likely to terminate than expected and were called as potential Psi sites. In addition, we expect biologically meaningful sites to show reproducibility across samples, thus we required that a potential Psi site was called in at least two *w*^*1118*^ replicates across the two independent experiments (details in Materials and Methods). Finally, we categorized the Psi sites according to high reproducibility (identified in 6-7 paired replicates), intermediate reproducibility (identified in 4-5 paired replicates), or low reproducibility (identified in 2-3 paired replicates).

According to these criteria, we identified 735 potential Psi sites on cellular rRNA which survived the polyA enrichment of our samples (**Table S3 in File S1**). Although this level of Psi on rRNA vastly exceeds the number of previously identified sites, 532 of the sites (72.4%) were highly reproducible. These highly reproducible sites on cellular rRNA are distributed across rRNA classes including 288 Psi sites in 28S, 16 sites in 5.8S, as well as 228 sites in 18S rRNAs (**Table S3 in File S1**). In *Drosophila melanogaster*, only 57 Psi sites on the large subunit (LSU) of cytoplasmic rRNA have been previously reported using the canonical primer-extension based pseudouridine detection method (Ofengand and Bakin 1997) (“Ofengand site” indicated by gray vertical lines in **Figure S2)**. Using our method of Psi calling, we have identified 53 of those known LSU sites (labeled as “Ofengand site” in Column H in **Table S3 in File S1**). Notably, 49/53 of those known sites are highly reproducible Psi sites in our analysis (**Table S3 in File S1**). In **Figure 2**, the cytoplasmic rRNA Psi sites are mapped onto the 3D based secondary structure of LSU rRNA of *Drosophila melanogaster* (RiboVision version 1.15+, (Bernier et al. 2014)). As there is no obvious clustering of the sites, additional analyses will be required to understand how the patterns observed relate to ribosomal function.

**Figure 2.**
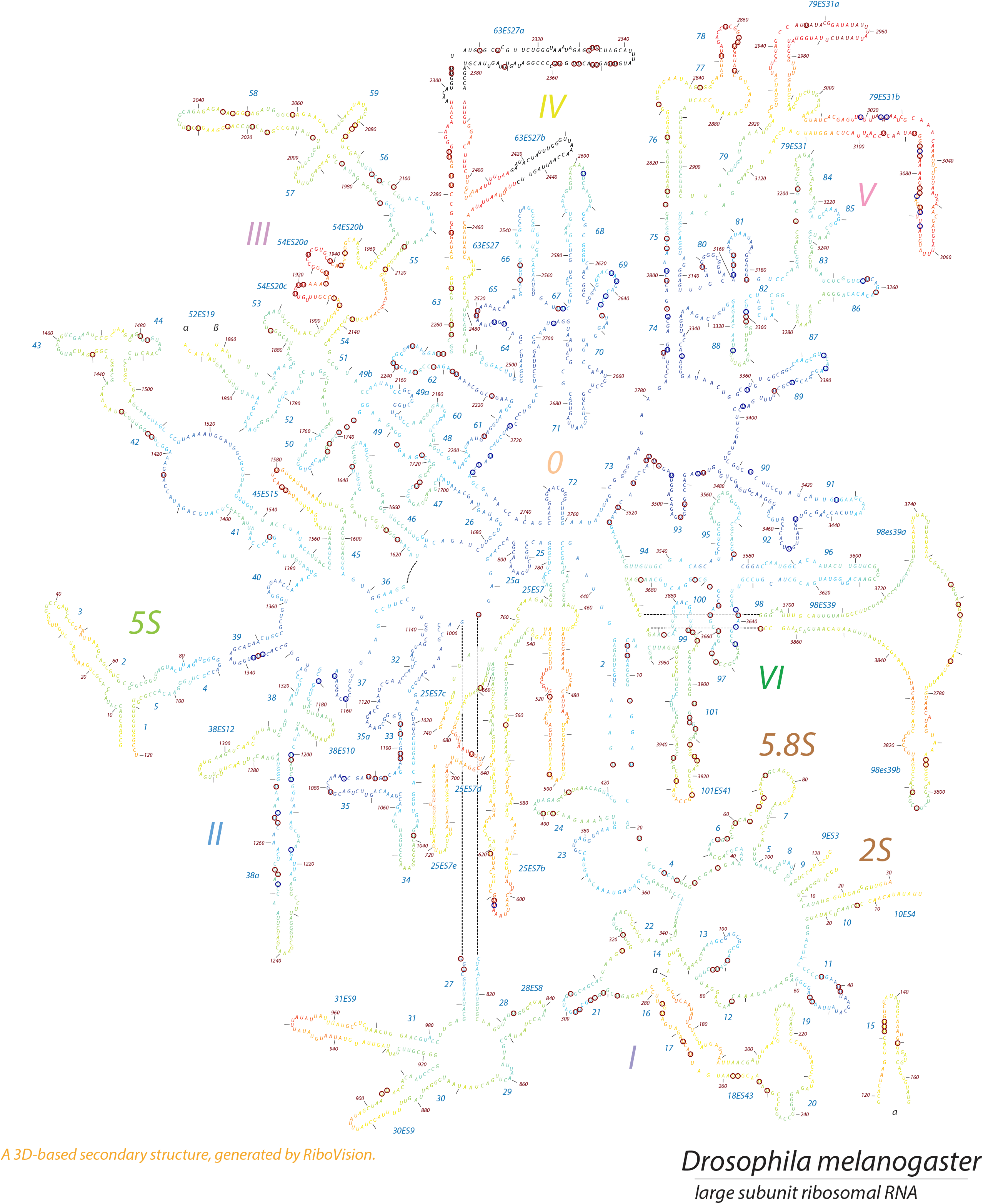
Location of pseudouridine modification sites (circled U) on cytoplasmic large subunit rRNA secondary structure in *Drosophila melanogaster*. Psi residues are circled in red for high reproducible sites and in blue for previously identified Offengand sites. The residues are color coded according to the “fine grained onion” scheme in which close proximity to the peptidyl transferase center is indicated by cooler colors (i.e. blue) and sites far from the center are indicated by warmer colors.

In addition to the 735 potential Psi sites identified on cytoplasmic rRNA, we also found 125 sites on mitochondrial rRNA, among which 57 highly reproducible sites were identified on the mitochondrial large ribosomal RNA (lrRNA) (**Table S3 in File S1**). Previously studies have identified a single pseudouridylation site in human mitochondrial 16S rRNA (Psi1397) which is essential for its stability and assembly into the mitochondrial ribosomes (Antonicka et al. 2017). Our results suggest a more widespread occurrence of Psi sites on mitochondrial rRNA in *D. melanogaster*.

### Psi-seq identified predicted and novel Psi in snRNAs, snoRNAs, tRNAs and other ncRNAs

Like the ribosome, the spliceosome is an important catalytic ribonucleoprotein machine. Numerous Psi sites have been reported in the spliceosomal snRNAs U1, U2, U4, U5, U6, U12, U4atac and U6atac (Yu et al. 2011; Adachi and Yu 2014). Although our protocol relied on polyA enriched RNA, we identified potential pseudouridine modifications on four spliceosomal snRNAs, including 21 on U1, 11 on U2, 2 on U5 and 5 on U6 (**Table S4 in File S1**). Six of these (Psi5, Psi6, Psi22 on U1, Psi44, Psi45, Psi55 on U2) were previously reported or predicted in Drosophila snRNAs (Deryusheva and Gall 2009; Huang et al. 2005). Additional sites (i.e. Psi46 on U5 and Psi40 on U6) were reported at the corresponding locations in human (Karijolich and Yu 2010) but have not been previously identified in Drosophila spliceosomal snRNA.

7SK is another important and highly conserved snRNA. 7SK is pseudouridylated at position U250 by the DKC1-box H/ACA RNP in human and this pseudouridylation was shown to be critical to stabilize 7SK snRNP and its function as a critical regulator of the homeostasis and activity of P*-*TEFb, a key regulator of RNA polymerase (pol) II transcription which stimulates the elongation phase (Zhao et al. 2016). This modification was identified from previous Psi-seq study (Carlile et al. 2014). The 7SK snRNA homolog and a similar P-TEFb control system function in *Drosophila* (Nguyen et al. 2012; Gruber et al. 2008). Our pseudo-seq analysis identified a single nucleotide pseudouridylation modification in d7SK snRNA, at U269 (**Table S4 in File S1**).

We found 10 Psi sites on the U3 RNA, a box C/D snoRNA required for rRNA processing. All the sites are distinct from the four Psi sites found on human U3 using mass spectrometry (Yamaki et al. 2020). There is also a single Psi site identified on snoRNA:Psi 28S-3342. Another 58 sites are localized on other ncRNAs, including 47 on 7SL, an abundant cytoplasmic RNA which functions in protein secretion as a component of the signal recognition particle (SSR). Previously only one Psi site (Psi211) has been reported on human 7SL. There are 11 sites on CR40469, an uncharacterized lncRNAs (**Table S4 in File S1**).

Our data indicates 8 Psi sites in Drosophila tRNAs (**Table S4 in File S1**), 3 of which (Psi13 and Psi54 of Glu-TTC and Psi27 of Val-CAC) were previously mapped on tRNAs of *D. melanogaster* (Machnicka et al. 2014).

All the Psi sites found on the non-coding RNAs, except for those on cytoplasmic and mitochondrial rRNAs in **Table S3 in File S1**, are listed in **Table S4 in File S1** and can be sorted by the RNA biotype (column F). Note that more non-coding RNA sites are likely to be identified using targeted approaches that do not rely on polyA enrichment.

### Psi modifications of mRNA transcripts

It was previously unknown if Psi occurs on *Drosophila* mRNAs. Our analysis identified 1147 mRNA Psi sites including 287 highly reproducible sites, 489 sites with intermediate reproducibility and 371 with low reproducibility (**Figure 3A, Table S5 in File S1**). Interestingly, a large proportion of pseudouridylated mRNAs have multiple sites identified in their transcripts (**Figure 3C**). Indeed, the 1147 Psi sites are localized on the transcripts of only 165 genes with transcripts from some loci containing more than 100 sites (**Figure 3C, Figure 3D, Table S5 in File S1**).

**Figure 3.**
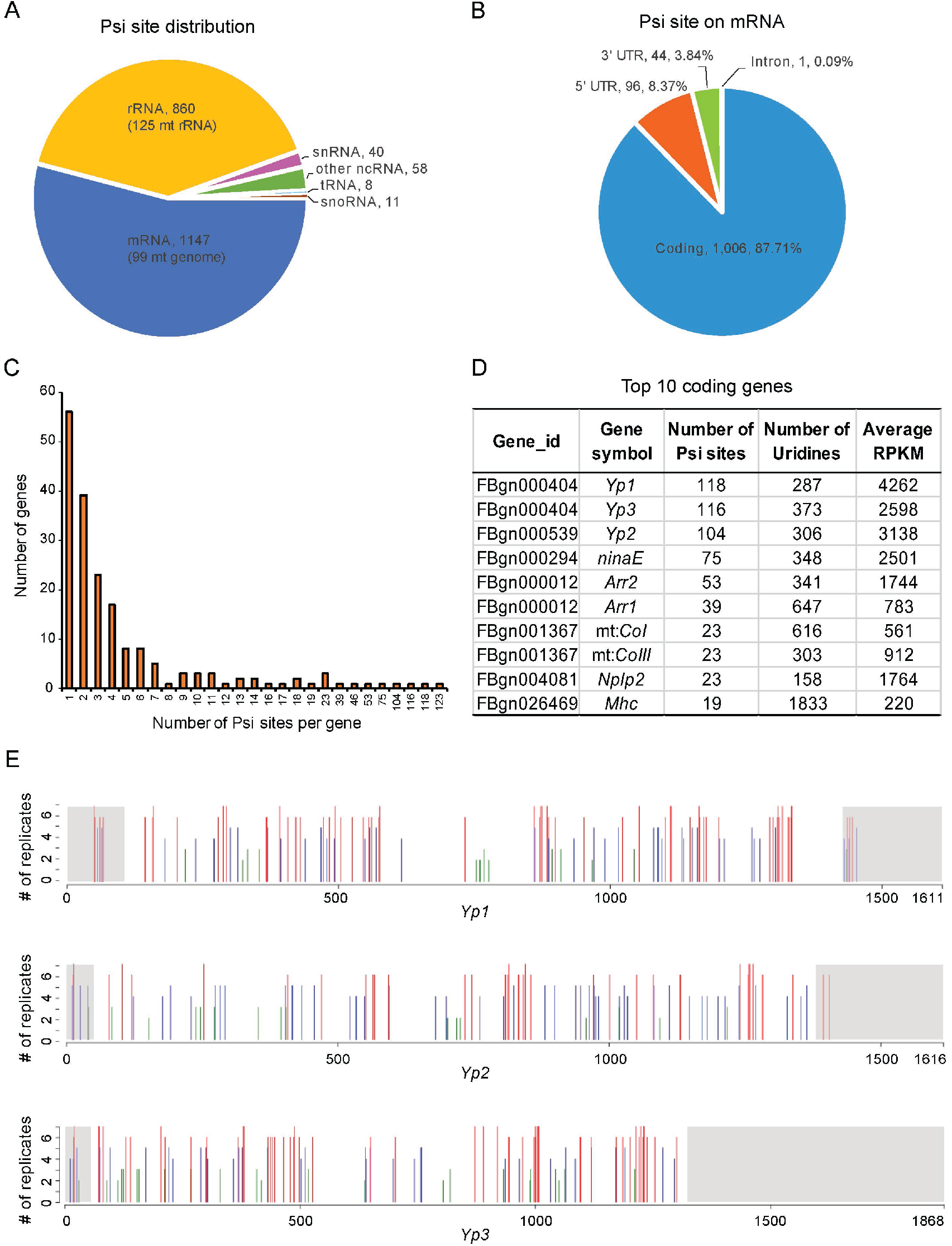
Features of pseudouridine sites identified from *Drosophila melanogaster* transcriptome. **A**. Psi site distribution according to RNA class (mRNA, rRNA, tRNA, snRNA, snoRNA and other ncRNA). **B**. Distribution of Psi sites on mRNA transcript regions (5’-UTR, coding, 3’-UTR and intron). **C**. Distribution of number of Psi sites identified per gene. **D**. The 10 coding genes with the highest number of Psi sites. For each gene, the numbers of Psi sites and Uridines in the sequence are listed along with the normalized gene expression represented as average RPKM of the *w1118* CMC libraries (n=7). **E**. Psi sites on yolk protein mRNAs. The locations of Psi sites identified on mRNAs for yolk protein *Yp1, Yp2*, and *Yp3* are indicated with bars above the schematic of the transcripts. UTRs are marked with gray shading whereas coding regions are not shaded. The height of the bars indicated by the Y axis represents the number of *w*^*1118*^ replicates a Psi site is identified from (n=2∼7) while red, blue and green color of the lines represents Psi sites of high, intermediate, and low reproducibility, respectively.

In *Drosophila*, the ribosomal large subunit (RpL, 60S) consists of three rRNAs (2S, 5.8S, 28S) and 47 ribosomal proteins whereas the small ribosomal subunit (RpS, 40S) consists of one ribosomal RNA (18S) and 33 ribosomal proteins (Anger et al. 2013). Our analysis identified 202 Psi sites in mRNAs encoding 35 of 47 RpL proteins and 23 of 33 RpS proteins. As with other mRNAs, most mRNAs encoding RPs contain at least two Psi sites. Although the vast majority the ribosomal proteins are pseudouridylated, we found no Psi modifications on the mitochondrial ribosomal protein mRNAs even though these were abundantly expressed in our RNA samples. Altogether, the 202 Psi sites on ribosomal protein RNAs accounted for 17.6% of total pseudouridines identified on mRNAs. In addition, if adding together the 735 Psi sites on the cytoplasmic rRNAs and 202 sites on various ribosomal protein mRNAs, nearly 44% of the Psi sites that we have identified are related to the production of cytoplasmic ribosomes.

Among the mRNA Psi sites, a total of 99 Psi sites were found on protein-coding genes of the mitochondrial genome (**Figure 3A, Table S5 in File S1**). Eleven of the 13 protein encoding mRNAs in the mitochondrial genome were pseudouridylated. Considering the 125 Psi sites found on the mitochondrial rRNA (**Table S3 in File S1**), we found a total of 224 Psi sites on the RNAs encoded from mitochondrial genome (10.5% of the total sites), thus indicating that RNAs from mitochondrial genome are significantly pseudouridylated in *Drosophila*.

As noted above, many of the mRNA transcripts have multiple Psi sites. Interestingly, Psi sites are found across functionally related mRNA transcripts. For instance, the mRNAs encoding the yolk proteins, which are known to be highly expressed in the fat bodies of adult female head (Fujii and Amrein 2002), contain the highest number of pseudouridine modification of the female head transcriptome. A total of 338 Psi sites (25.5% of total mRNA Psi sites) were identified on mRNAs encoding the three yolk proteins (Yp1, Yp2, and Yp3). Each of the yolk protein mRNAs harbors many Psi sites (118 sites on *Yp1*, 104 on *Yp2* and 116 sites on *Yp3*) (**Figure 3E, Table S5 in File S1**). Notably, the majority of the Psi sites are localized in the coding region and are almost absent from the 3’ UTR region of the transcripts, which confirms the specificity of our Psi-seq analysis and strongly suggests for an important function of pseudouridylation in the coding region for the yolk protein transcripts.

Transcripts of multiple genes involved in sensory function in the nervous system are also marked with Psi modifications. A notable example is *ninaE*, the structural gene encoding the *Drosophila* Rhodopsin-1, with 75 Psi sites. Indeed, a number of transcripts bearing multiple Psi sites play various roles in photoreceptor function, including other *rhodopsins* (*Rh*) (*Rh3*: 2 Psi sites, *Rh4*: 2 Psi sites, *Rh6*: 2 Psi sites), *retinin* (18 Psi sites), and *Arrestins* (*Arr1*: 39 Psi sites, *Arr2*: 53 Psi sites). Several *Odorant-binding protein* mRNAs, which function in the sensory perception of smell, also have multiple Psi sites (*Obp19d*: 3 Psi sites, *Obp44a*: 12, *Obp56d*: 3, *Obp56e*: 2, *Obp99c*: 1). It is interesting that mRNAs encoding two potential neuropeptide genes *Nplp2* and *Nplp3* contain 28 and 23 Psi sites, respectively (**Table S5 in File S1**).

Previously, an N3-CMC-enriched pseudouridine sequencing identified pseudouridylation at U519 of the human elongation factor *eEF1A1* mRNA which encodes the main protein that delivers aminoacyl-tRNAs to the ribosome. In fact, *eEF1A1* is the first experimentally validated human mRNA psi site (Li et al. 2015). From our Psi-seq data, sites were identified on mRNA for two of the *Drosophila* translation elongation factors (*eEF1α1* with four sites and *eEF1γ* with one site) (**Table S5 in File S1**).

The majority of Psi sites (1006 sites, 87.7%) found on mRNAs were within the coding region. The 5’ untranslated regions (UTRs) had only 98 (8.5%) sites and 61 sites (5.3%) were in 3’ UTRs (**Figure 3B**). In coding regions, Psi sites were more commonly found in the second position of the codon relative to the first or third positions. For nuclear-encoded mRNA, the most highly pseudouridylated codons are the CΨG and UΨG for Leucine, followed by GCΨ for Alanine and ΨGG for Tryptophan (**Table 1**). Interestingly, the pattern of pseudouridylation of codons in mitochondrial genome transcripts appeared to be distinct from that of nuclear genome transcripts, with the most frequent Psi-containing codons encoding for Phenylalanine (F) (i.e. UUΨ and UΨU), Isoleucine (I) (i.e. AΨT), as well as Leucine (but in the mitochondrial case, ΨTA was the most prominent codon for Leucine) (**Table 1**).

**Table 1.**
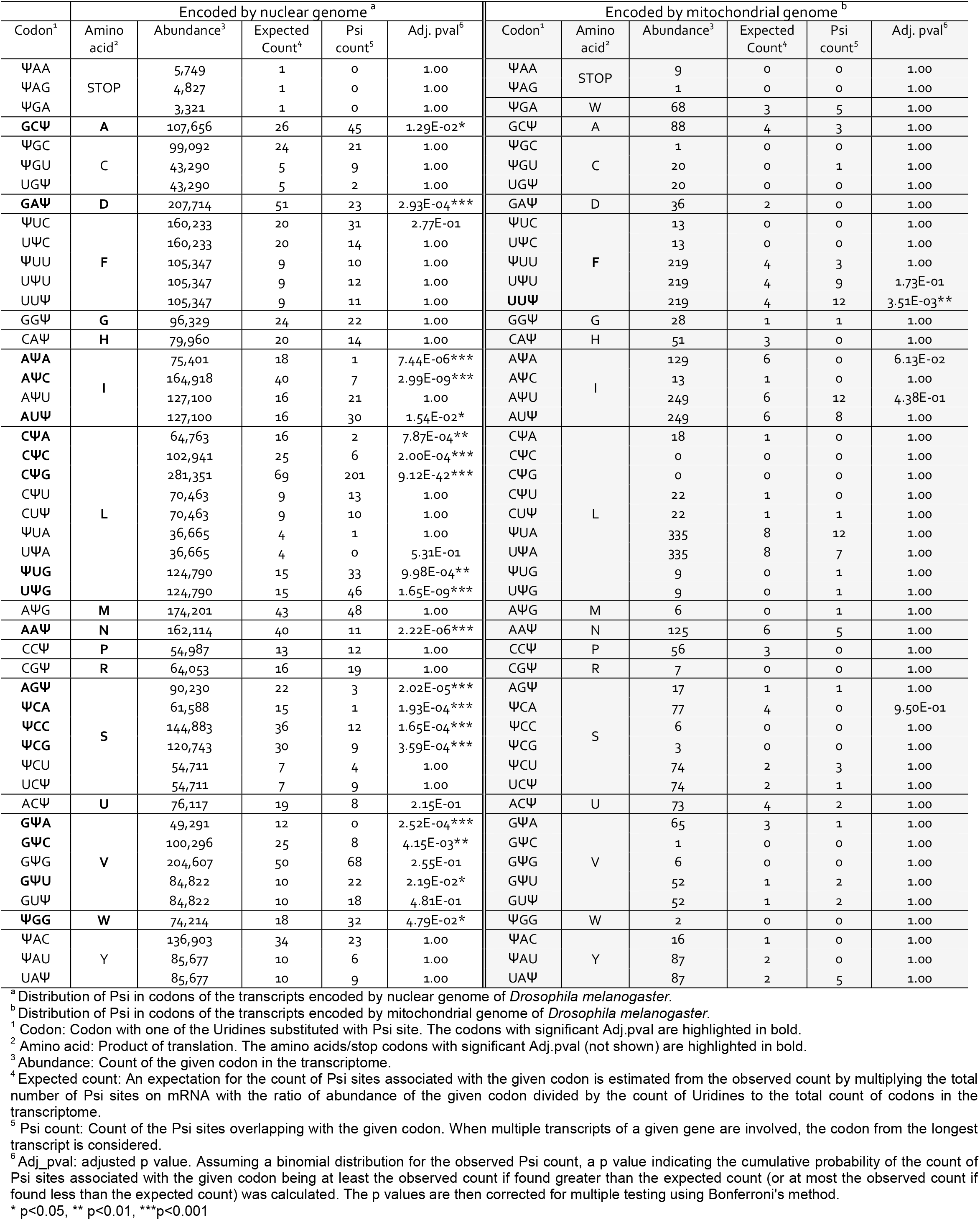
Distribution of Psi (Ψ) sites on nuclear (left) or mitochondrial (right, shaded) genome encoded codons and amino acids in *Drosophila melanogaster*.

### Pseudouridylation of the UGA codon in Drosophila mitochondria

A previous study proposed that Psi in mRNA stop codons may result in the suppression of translation termination and enhancement of translational read-through *in vitro* protein synthesis (Karijolich and Yu 2011). We were very interested in this as a set of transcripts in the *Drosophila* transcriptome are predicted to undergo stop codon suppression (Jungreis et al. 2011). However, our analysis failed to identify any Psi sites on stop codons of nuclear encoded genes (**Table 1, Table S5 in File S1**). This result suggests that pseudouridylation is unlikely to be an important underlying mechanism for stop-codon readthrough in *Drosophila*.

However, we did find Psi sites present on the canonical “stop codon” UGA in mitochondrial encoded transcripts. Out of the 68 total UGAs in mitochondrial mRNA, 5 were called as ΨGA from our analysis. This was in contrast to 3,321 UGAs that were present among the nuclear mRNA that did not contain a signal for ΨGA. The frequency of ΨGA codons in mitochondrial RNA was significantly higher than in nuclear encoded RNA (Fishers exact test p-value = 2.81e-9). Importantly, the UGA codon in mitochondria of insects and in vertebrates does not encode a stop, but instead codes for tryptophan. Our analysis suggests that this codon, at least in some cases, is actually ΨGA.

### mRNA pseudouridylation is correlated with abundance

The yolk protein mRNAs are among the most highly expressed transcripts in our female fly head libraries and these were the most highly pseudouridylated genes. mRNAs encoding ribosomal proteins are also abundantly expressed and were among the classes of genes with a high likelihood of Psi modification in their transcripts. Thus, we explored the relationship between Psi occurrence and RNA expression level in CMC library (**Figure 4A)**. Interestingly, for mRNAs there appeared to be a strong positive correlation between mRNA expression level and the number of Psi sites in a particular transcript. The majority of the Psi sites are identified on the transcripts of the genes which are abundantly expressed (**Figure 4B**). However, this was unlikely to be caused by a limitation coverage in calling sites because we found no correlation between the read coverage and the presence of Psi sites for the non-coding RNAs, such as snRNA, snoRNA, tRNA and other nc-RNA (**Figure 4A**). Our results don’t allow us to infer a cause and effect for the frequency of Psi on *Drosophila* mRNAs. Highly expressed mRNAs may be more stable due to the presence of Psi modification, or they may be more likely to be pseudouridylated because they are more highly expressed, or both factors may contribute to varying degrees.

**Figure 4.**
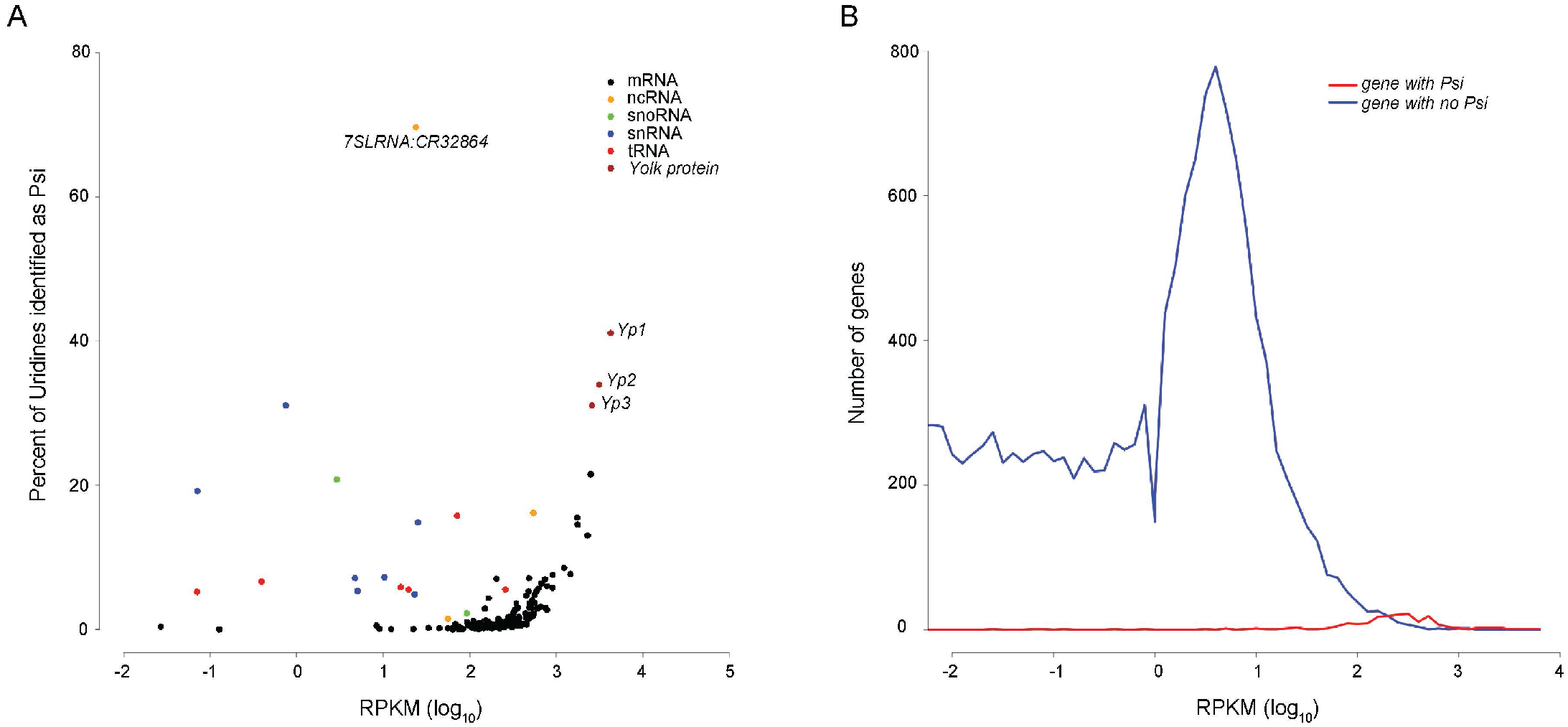
Correlation between gene expression and Psi site identification. **A**. A scatter plot of gene expression *versus* proportion of Psi site across pseudouridylated transcripts. The gene expression is represented as the average of log10 (RPKM) in *w*^*1118*^ CMC libraries (n=7) and the proportion of Psi sites is measured by the percentage of Psi sites called on the total number of Uridines in various categories of RNA species. The individual data points representing the yolk proteins and 7SL RNA are labeled. **B**. A plot of gene expression *versus* the number of genes with (red) or without (blue) Psi sites identified on their transcripts.

### Transcriptome-wide identification of potential pseudouridylation targets of RluA-2

Previously, the putative pseudouridine synthase RluA-2 was found to be broadly expressed in *Drosophila* and loss of function mutation in *RluA-2* was found to cause hypersensitivity in larval pain pathways (Song et al. 2020). Thus, we sought to identify potential targets for RluA-2 by comparing the pseudouridine profiles of *RluA-2* mutant mRNA with *w*^*1118*^ control mRNA. A total of 423 Psi sites were not detected in any of the four *RluA-2* mutant libraries. These missing sites therefore represent potential targets of RluA-2 (**Table S6 in File S1**). The candidate RluA-2 target sites include 74 pseudouridylation sites on cytoplasmic and mitochondrial rRNAs (63 cellular and 11 mitochondrial), 5 on snRNAs, 5 on tRNA, 9 on other ncRNAs and 328 sites on mRNAs (**Figure 5**). The mRNA Psi sites include 65 of the sites encoding for components of ribosomal proteins and 42 mitochondrial genome-encoded transcripts. Among the 165 genes where the pseudouridylation sites were identified on *w*^*1118*^ transcripts (**Table S5 in File S1**), 115 genes were missing Psi modifications in the *RluA-2* mutant. That the sites were missing in various classes of RNA suggests that the RluA-2 enzyme acts on a wide range of substrates (**Table S6 in File S1**).

**Figure 5.**
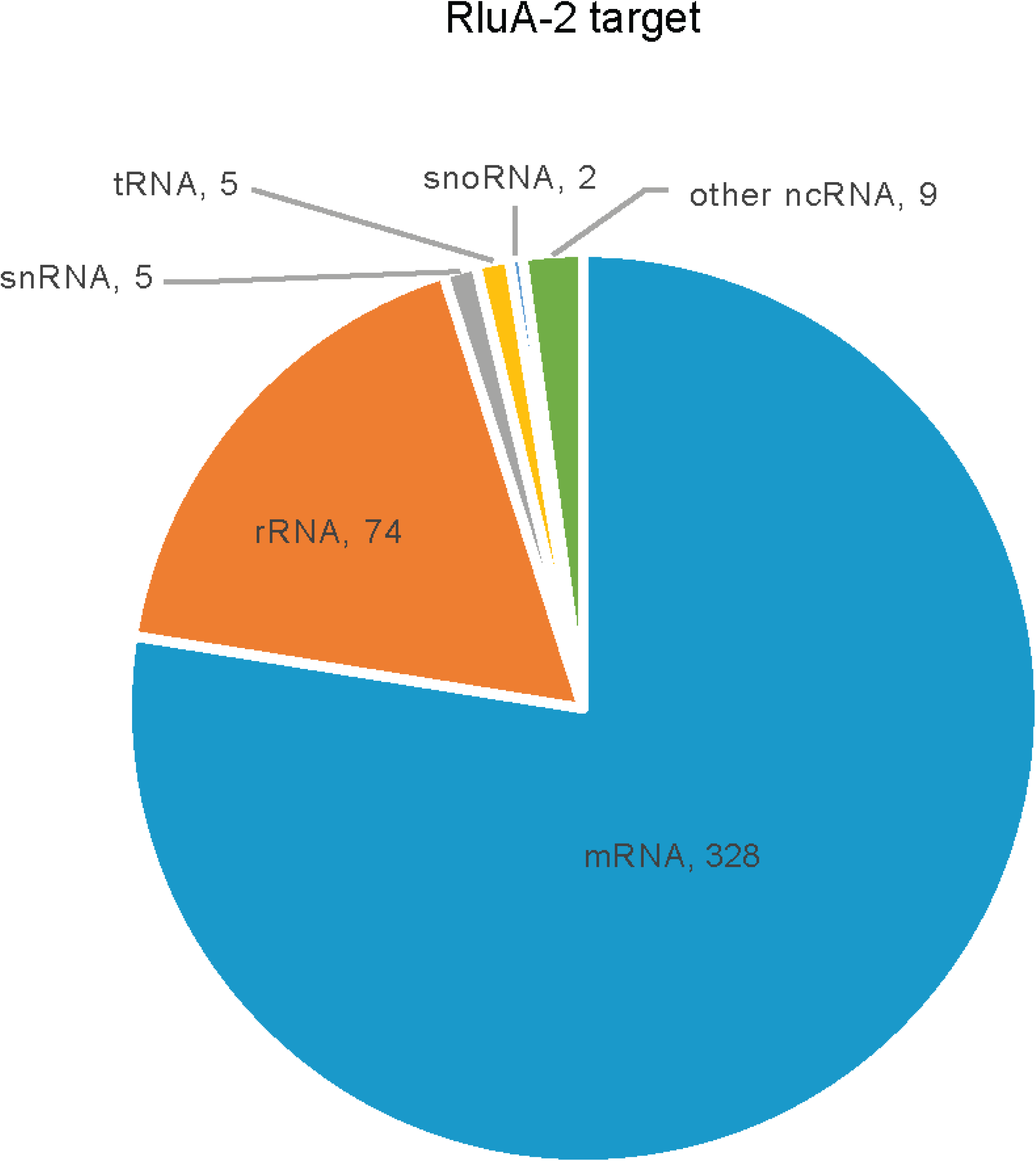
Pie chart showing the distribution of number of RluA-2-dependent Psi sites in different categories of RNA species (mRNA, rRNA, tRNA, snRNA, snoRNA and other ncRNA). Those potential RluA-2 target sites are missing from *RluA-2* mutant compared to the *w*^*1118*^.

To determine if RluA-2 recognizes a particular sequence motif we ran the target sites through a motif finding algorithm. Other than a dominant G downstream of the pseudouridine site there was not a significant sequence logo around the modification site (**Figure S3**). This finding suggests that RluA-2 enzyme may not recognize a primary sequence motif in site selectivity and may indicate that secondary structure of RNAs is critical for RluA-2 target recognition (as has been found for other pseudouridine synthases (Purchal et al. 2022)).

### Loss of RluA-2 results in differential expression in translation-related genes and non-coding RNAs

To investigate the effects of *RluA-2* on gene expression we compared the transcriptome of *RluA-2* mutant to the control genetic background *isow*^1118^ in mock treated libraries of experiment 2. The Euclidean distance of the libraries demonstrated a clear clustering of samples based on genotype **(Figure S4)**. A total of 2214 genes showed significant change in their expression levels in the *RluA-2* vs. *isow*^1118^ samples (FDR<0.05), including 1048 that were upregulated and 1166 that were downregulated in the mutant (**Figure 6A, Table S7 in File S1**). The vast majority of differentially expressed genes (94.8%) were mRNA-coding genes, which included 994 of the upregulated and 1104 of the downregulated genes. Among the upregulated gene set, we found 35 RpL genes and 22 RpS genes, three elongation factors and 11 initiation factors (**Figure 6B, Table S7 in File S1**). It is notable that *Drosophila* Myc (dMyc), which is a major regulator of genes involved in ribosome biogenesis and translation (Gallant 2013), as well as its binding partner dMax, were both upregulated in *RluA-2*. Consistent with these observations, a gene set enrichment analysis of the differentially expressed genes showed significant enrichment in the ribosome (FDR < 0.0001) and ribosome biogenesis (FDR = 0.154) (**Figure 6C**). These results suggest a potential impact on the translation and translational machinery that is caused by loss of RluA-2.

**Figure 6.**
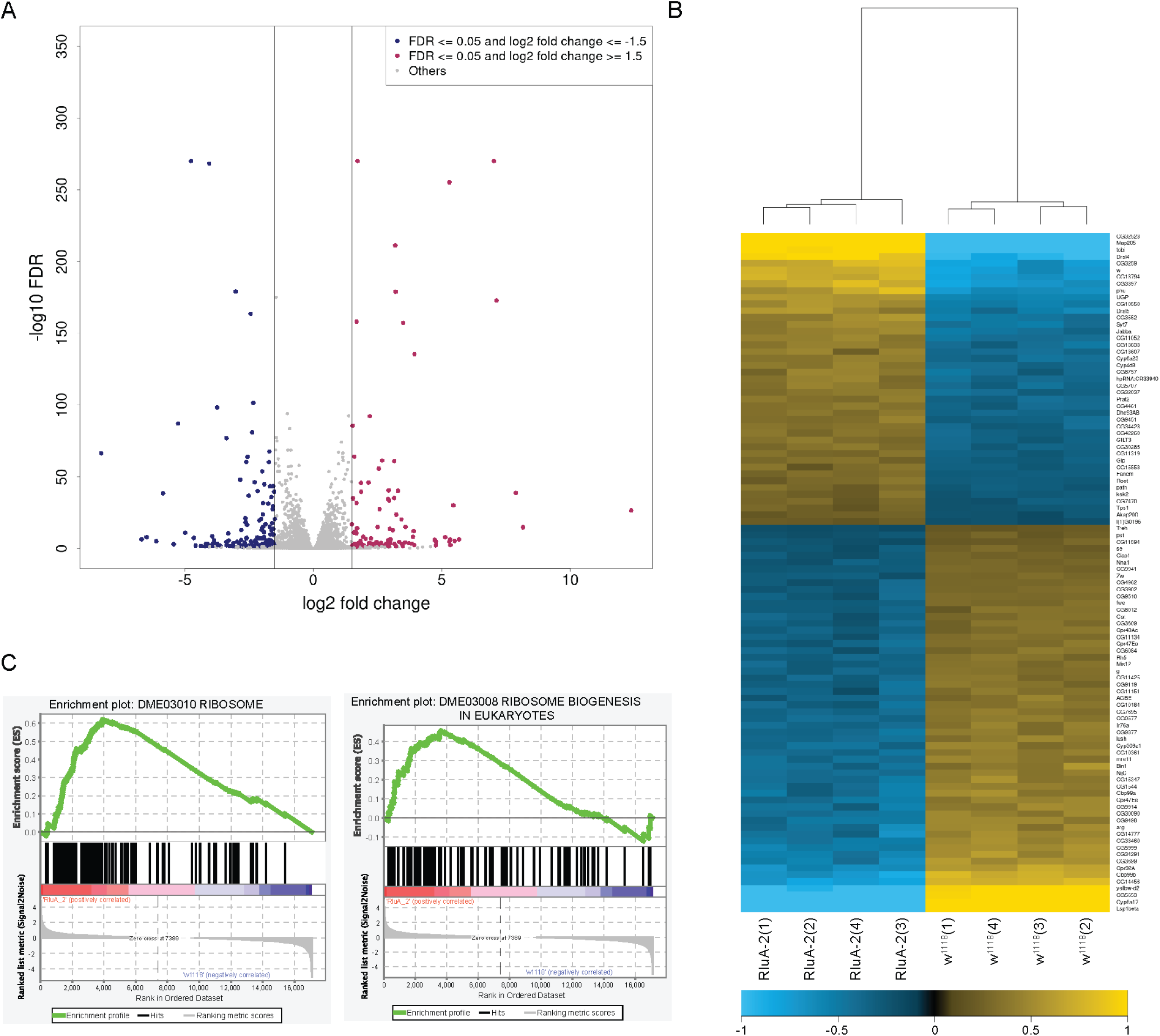
Effect of loss of RluA-2 activity on gene expression in the female *Drosophila* head transcriptome. **A**. Volcano plot showing the differentially expressed genes in the mock Psi-seq libraries of *RluA-2* vs *w*^*1118*^. The log_2_ fold change indicated the mean expression level for each gene. Each data point represents an individual gene. Red data points indicate upregulated genes with log_2_fold change>=1.5, the blue data points indicate downregulated genes with log_2_fold change<=1.5 in *RluA-2* vs *w*^1118^ (FDR<=0.05) and grey data points represent all the other genes. **B**. A hierarchically clustered heatmap showing the expression pattern of the top 100 significantly differentially expressed genes between *RluA-2* vs *w*^1118^. Gene expression values quantified as the counts of reads mapping to the exon regions were transformed by applying DESeq2’s regularized-logarithm transformation. The rows representing the genes were sorted in descending order of their average expression among the *RluA-2* libraries. The columns representing the individual libraries were hierarchically clustered based on the gene expression. The color density on the heatmap indicates the extent of deviation for each gene in each replicate library from the gene’s average expression across all libraries. Yellow and blue represent up- and down-regulated expression in *RluA-2*, respectively. **C**. Gene set enrichment plots show that the Gene Ontology categories are enriched for the Ribosome (left) and Ribosome Biogenesis in Eukaryotes (right) in the *RluA-2* mutants. The y-axis in the top portion represents enrichment score (ES) and on the x-axis are genes (vertical black lines) represented in genes sets. The green line connects points of ES and genes. ES is the maximum deviation from zero as calculated for each gene going down the ranked list, and represents the degree of over-representation of a gene set at the top or the bottom of the ranked gene list. The middle portion shows where the members of the gene set appear in the ranked list of genes. The colored band represents the degree of correlation of genes with *RluA-2* (red for positive and blue for negative correlation). The ranking metric at the bottom portion shows a gene’s correlation with *RluA-2* (positive value) or *w*^1118^ (negative value). The significantly enriched gene sets are DME03010 RIBOSOME (p value = 0.000; FDR = 0.000) and DME03008 RIBOSOME BIOGENESIS IN EUKARYOTES (p value = 0.000; FDR = 0.154).

Notably, 77 non-coding RNAs (non-protein coding anti-sense RNAs and long non-coding RNAs) were either upregulated (47) or downregulated (30) and several snoRNAs responsible for guiding pseudouridylation or methylation on 18S and 28S rRNAs were among the differentially regulated RNAs (**Table S7 in File S1**).

We investigated impact of loss of RluA-2 on the other eight putative pseudouridine synthases. Expression of *RluA-1*, the paralog of *RluA-2*, was slightly decreased which may be a consequence of loss of regulatory DNA in the *RluA-2* deletion. In contrast, *mfl* the human *Dyskerin* homolog, and *CG3709* (encoding the putative *Drosophila* Pus10), were upregulated although the fold change in expression was also slight (**Table S7 in File S1**). The expression of the other putative pseudouridine synthases is either not changed (*CG3045, CG4159*/*PUS1, CG34140*) or not detected in female head (*CG7849* and *CG6745/PUS7*).

Finally, we investigated whether there might be a correlation between RluA-2 dependent pseudouridylation and gene expression. If such a correlation exists, transcripts that lose sites in the mutant might show altered transcript levels. Interestingly, 67 genes with missing Psi sites in the *RluA-2* mutant also show differential expression (**Table S8 in File S1**). Among the RluA-2 targets, ribosomal protein component genes are also differentially expressed (18/21 *RpL* and 9/15 *RpS* genes). The differentially expressed mRNAs encoding ribosomal proteins all showed upregulation in the *RluA-2* mutant. A similar trend of upregulation was also observed in other RluA-2 target genes, including yolk protein mRNAs (*Yp1, Yp2, Yp3*). However, some transcripts missing Psi sites were found to be reduced in expression (i.e. *Nplp3, Obp19d, Obp44a*). The vast majority of the differentially expressed genes did not contain RluA-2 dependent Psi sites (2147, 97.0%), suggesting the majority of the differentially expressed genes do not change expression levels as a consequence of a missing pseudouridylation. This may indicate that *RluA-2* can directly affect RNA levels independently of its pseudouridine synthase activity, or the gene expression alterations may be a downstream or indirect consequence of a lack of *RluA-2* activity. Future studies investigating the RNAs that are directly bound by RluA-2 will be necessary to disentangle these alternatives.

## Discussion

For over a century studies of *Drosophila melanogaster* have led to countless insights into the underlying principles of genetics and molecular biology including several studies of pseudouridylation of RNA. Nevertheless, next generation sequencing has yet to be applied to thoroughly characterize *Drosophila* Psi. We have described here the first attempt at characterizing a *D. melanogaster* pseudouridine epitranscriptome. Our optimized library construction and bioinformatics workflow has allowed us to robustly detect previously known sites as well as to uncover a set of previously unknown Psi sites that are reproducible across biological replicates. Although our analysis has been limited to a single stage of the fly life cycle, and a single sex, our findings provide interesting insights for future studies.

Perhaps the most striking finding of our study is the widespread presence of Psi in the RNAs that encode for the many components of the cellular ribosome. Over 500 highly reproducible sites were found in rRNA which is significantly more than the number of previously identified sites. The vastly increased number of candidate rRNA sites that we have detected is likely due to the statistical power that is provided with next generation sequencing methods. A caveat to consider, is that our methods do not allow for the explicit determination of the proportion of pseudouridylated bases that occur at individual sites. Even for highly reproducible sites, it is likely that these sites occur on only a fraction of rRNA molecules and additional studies will be needed for quantitative determination of site frequency. This could potentially be performed with recently described HydraPsiSeq methodology (Marchand et al. 2020). Alternatively, nanopore sequencing approaches would potentially allow for determination of Psi sites that occur together on individual rRNA molecules (Begik et al. 2021).

In addition to sites on rRNA, we found that the vast majority of mRNAs encoding ribosomal proteins were pseudouridylated. Although the function of the Psi sites on *Drosophila* mRNAs also awaits further analysis, an interesting possibility is that codons containing Psi could favor the incorporation of non-cognate amino acids during mRNA translation (Eyler et al. 2019). We found 160 Psi containing codons across mRNAs encoding for 58 ribosomal proteins. If each of these codons facilitates some degree of non-cognate amino acid incorporation (Eyler et al. 2019), this would allow for vast complexity of ribosomal protein isoforms. We hypothesize that such a diversity of ribosomal protein subunits might allow for robustness in ribosomal function across a variety of environmental conditions which could be advantageous for a poikilotherm such as *Drosophila*. Alternatively, ribosomal protein diversity could contribute to cell type specific functions of the ribosome, or facilitate translation of particular mRNAs as previously described for other forms of ribosome heterogeneity (Shi et al. 2017; Genuth and Barna 2018; Hopes et al. 2022).

The most highly pseudouridylated mRNAs (*Yp1, Yp2*, and *Yp3*) encode for the Yolk proteins. These transcripts are expressed in fat bodies of the female head which is one of the places where yolk protein biogenesis is thought to occur (Fujii and Amrein 2002). An important aspect of fly vitellogenesis involves synthesis and secretion of the yolk proteins from fat bodies into the hemolymph circulation for transport to the ovaries (Hames and Bownes 1978). High levels of pseudouridylation, especially in the coding region of the Yp transcripts, may facilitate message stability, translation, or possibly plays a role in targeting the mRNA to secretory pathways. Consistent with the latter possibility, the RNA with the highest percentage of Psi was the 7SL RNA which is a component of the Signal Recognition Particle, the key factor in guiding mRNAs to the secretory compartments of the cell. Other mRNAs for secreted and transmembrane proteins that identified in our analysis include rhodopsins (*ninaE, rh4, rh6*), *trp*, and odorant binding proteins (*Obp44a, Obp56d, Obp56e, Obp99c, Obp19D*).

We found that transcripts expressed from the mitochondrial genome (*lrRNA, srRNA, CoI, CoII, CoIII, AtpAse6, AtpAse8, ND2, ND5, Cyt-b)* and some nuclear encoded mitochondrial genes (*ATPsynB, ATPsynbeta, ATPsynC, ATPsynF, Cyt-c-p, COX6B, COX7C, UQCR-C2, COX5A*, and *porin*) were pseudouridylated. Interestingly, the opal (UGA) cellular stop codon was pseudouridylated in 5 cases on mitochondrial transcripts and we propose that this may facilitate incorporation of tryptophan at ΨGA codon as part of the mitochondrial genetic code. Indeed, our finding that the ΨGA codon is specific to mitochondrially encoded RNAs provides evidence in support of specificity of the Psi sites that we have identified. If our method were to identify spurious sites, we would expect to find ΨGA codons in both nuclear and mitochondrially encoded genes. Other potential Psi sites that were absent from our analysis are further evidence of specificity (**Table 1**). Notably, we found no instances for Psi on the amber (UAG) or ochre (UAA) stop codons. Indeed, many codons were detected at lower levels than expected if Psi were equally distributed across uridine nucleotides (**Table 1)**.

An interesting pattern is apparent for nuclear and mitochondrial encoded genes and their pseudouridylation of codons for Leucine. All of the seven instances for the TΨA codon (i.e. TTA Leucine) were found on mitochondrially encoded mRNAs. We also found evidence for ΨTA modification of the TTA codon, and in 12/13 cases this modification was identified on mitochondrial encoded transcripts. Interestingly, on nuclear encoded transcripts, the CUG codon for Leucine was the codon that was most enriched for Psi. As the CUG codon for Leucine is not found at all in mitochondrially encoded genes, all of the instances of CΨG codons are in nuclear encoded genes (as expected). Thus, in mitochondria there is an enrichment for Psi in the UUA codon for leucine but for nuclear encoded genes the enrichment is for the CUG codon for leucine. We hypothesize that this use of Psi by the cell may contribute to the appropriate translation of biased codons for Leucine on mitochondrial and cellular ribosomes respectively. We note that Leucine (along with Serine and Arginine) is unusual among amino acids in that it can be encoded by six codons (using the canonical bases A, C, U and G). If one considers that there are 11 possible Psi containing codons for Leucine, the total number of potential codons for Leucine expands to 17. The mechanistic importance of this level of diversity for Leucine codons poses an interesting puzzle for the future.

Finally, we found that 423 Psi sites were absent from the *RluA-2* mutant RNA and these sites are thus potential targets for the RluA-2 enzyme. The potential RNA targets of RluA-2 included rRNAs, mRNAs, snRNAs and tRNAs. Among mRNAs, notable targets for RluA-2 included *arrestin2 (Arr2)* mRNA which was missing 24/42 sites and *neuropeptide like protein 3 (NPLP3*) was missing 11/14 sites. These two targets stand out as potentially important in the previously described hypersensitive nociception phenotypes of *RluA-2* mutant larvae (Song et al. 2020). Ribosomal protein mRNA targets of RluA-2 are also of note as 65/160 sites on mRNAs encoding for ribosomal proteins were absent in the mutant background. Interestingly, many of these same ribosomal protein mRNAs showed upregulated expression in the mutant RNA libraries. We look forward to future studies that may further elaborate the impact of RluA-2 dependent pseudouridylation on neuronal and ribosomal function.

## Supplemental Materials

## Supplemental Figures

**Figure S1**. Optimization of CMCT treatment conditions for Psi-seq library preparations. The polyA-selected RNA was subjected to either fragmentation (“Fragmented before treatment”) or not (“Non-fragmented before treatment”) prior to the CMC/mock derivatization. The derivatization was carried out with CMC/mock treatment for 1hr or 18hr at 37°C (“1hr CMC/mock” or “18hr CMC/mock”), followed by alkaline treatment for 6hr at 37°C or 75°C (“37°C hydrolysis” or “75°C hydrolysis”). The Y axis represents the termination fold change while the X axis is the nucleotide position on the RNA spike-in sequence. The termination fold change from the only pseudouridine at position 43 on the RNA spike-in is marked with a red line whereas those from other sites are presented as black lines. The optimum condition producing the highest termination fold change for Psi site and lowest background is framed with a dashed red rectangle.

**Figure S2**. The termination fold change (y axis) distribution over the large subunit (LSU) of cytoplasmic rRNA in *Drosophila melanogaster* (x axis). The Psi sites identified are called with the threshold of appearing in 6 or all 7 *w*^*1118*^ libraries (high reproducible site, red), 4 or 5 libraries (intermediate reproducible site, blue), and 2 or 3 libraries (low reproducible site, green) from both sets of the experiments. The previously reported Psi sites on LSU identified by Ofengand et al (1997) were indicated by the gray vertical lines.

**Figure S3**. Consensus sequence for the RluA-2 target sites. The X axis is the +/-10 nucleotide sequence flanking the T site where the RNA bears the pseudouridylation modification (0 position). The Y axis represent the bits score. The overall probability of a nucleotide is represented by the height of the letter.

**Figure S4**. Euclidean distances of *w*^*1118*^ and *RluA-2* mutant mock libraries clearly demonstrate the clustering of samples based on the genotypes. Color key indicates level of similarity between libraries.

Supplemental File S1

**Table S1**. A summary of the Psi-seq libraries (Table S1) and an overview of contents included in supplemental tables (Table S2 - Table S8) for transcriptome-wide mapping of pseudouridylation and transcriptome expression analyses in *Drosophila melanogaster*.

**Table S2**. The quality features of the sequencing data used for Psi-seq and transcriptome expression analyses in *Drosophila melanogaster*.

**Table S3**. Psi sites identified using Psi-seq on cytoplasmic and mitochondrial rRNA in *Drosophila melanogaster*. Note the cytoplasmic LSU rRNA sites reported previously are labeled as “Ofengand site” in column H.

**Table S4**. Psi sites detected on other types of ncRNA with Psi-seq, including tRNA, snRNA, snoRNA, and other ncRNA in *Drosophila melanogaster*.

**Table S5**. Psi sites detected on protein-encoding mRNAs with Psi-seq in *Drosophila melanogaster*.

**Table S6**. Potential RluA-2 pseudouridylation targets in the adult head of *Drosophila melanogaster* with Psi-seq. Pseudouridylation sites from *RluA-2* (n=4) and *w*^*1118*^ (n=4) samples from the same set of experiment (Exp 2) were compared and those sites which are called in at least two libraries from both of the experiments (Exp 1 and Exp 2) in *w*^*1118*^ but absent from *RluA-2* were listed as the putative RluA-2 targets.

**Table S7**. Differentially expressed genes in *RluA-2* compared to the genetic background iso*w*^*1118*^ in the adult head of *Drosophila melanogaster*. The mock samples from each Psi-seq data set were used for differential expression analysis between the 4 replicates of *RluA-2* samples and 4 replicates of *w*^*1118*^ samples. Differentially expressed genes were detected using DeSeq2 with the threshold of adjusted *p* value with IHW correction <0.05.

**Table S8**: A list of differentially expressed genes between *RluA-2* and *w*^*1118*^ which overlap with genes containing RluA-2 target Psi sites.

## Materials and Methods

### Fly Strains and Raising Conditions

The isogenic *w*^*1118*^ strain used as the control genotype was obtained from Bloomington stock center. The mutant of *RluA-2*^*del-FRT*^ used in the study was a complete deletion of the locus that was generated and previously described (Song et al. 2020). All flies were reared on yeast, corn meal food in bottles or vials at an incubator with controlled temperature (25°C) and humidity (70%) on a 12h light/12h dark cycle.

### RNA extraction from adult head tissue

For Experiment 1, newly eclosed virgins from iso*w*^*1118*^ were collected twice every day (once in the morning and once in the afternoon) with less than 8 hours in between. For Experiment 2, newly eclosed virgins from iso*w*^*1118*^ and homozygous *RluA-2* ^*del-FRT*^ (Song et al. 2020) were collected in the similar way. The flies were put into vials with yeast and Bloomington recipe fly medium (20-30 flies in each vial) and aged for 1d in the incubator. The flies were then snap frozen in liquid nitrogen and kept at -80°C for 3 days until sufficient number of flies from each genotype had been collected. The frozen flies were then decapitated and 100 heads per replicate homogenized in TRIzol reagent (Invitrogen). Total RNA was prepared according to the manufacturers’ recommendations.

### In vitro transcription of RNA “spike-in”

The RNA spike-in sequence containing a single pseudouridine at position 43 as used previously (Schwartz et al. 2014) was *in vitro* transcribed from dsDNA template (synthesized as gBlocks gene fragment from IDT). *In vitro* transcription was performed in a volume of 20ul with MEGAshortscript T7 Transcription kit (AM1354, Invitrogen), using 75nmol each of GTP, CTP, ATP and Psi-TP (TriLink Biotechnologies). The RNA product was purified with MEGAclear Transcription Clean-Up kit (AM1908, Invitrogen) before adding into the Psi-seq libraries.

### CMCT treatment and RNA-seq library preparation

Polyadenylated RNA was enriched from total RNA using two rounds of Oligo(dT) dynabeads (Invitrogen) according to manufacturer’s protocol. To each polyA-RNA sample, spike-in RNA (∼3ug) was added and then the samples were split into two equal samples for CMCT or mock treatment. CMCT treatment was carried out essentially as described (Bakin and Ofengand 1993, 1995). However, when we used the exact conditions of prior studies (Schwartz et al. 2014), we failed to obtain a distinctive signal of termination fold change at the expected Psi position of our spiked in RNA (data not shown). Therefore, to optimize the CMC treatment and pseudouridylation detection efficiency for our Psi-seq, we tested the impact of several main factors affecting the efficiency of CMCT derivatization, including fragmentation before CMC treatment as used previously (Li et al. 2015; Carlile et al. 2014), alternative temperatures and durations of CMCT and alkaline treatment. We found that a 1hr CMCT treatment at 37°C on non-fragmented RNA combined with the 6 hours of alkaline hydrolysis at 37°C resulted in the highest termination fold change (>5) at the expected position in the spike-in and significantly reduced noise compared to a variety of other treatment conditions that we attempted (**Figure S1**). We thus implemented this optimized CMCT treatment conditions for our Psi-seq library preparation.

Specifically, pelleted polyA+ RNA was resuspended in 30ul 200mM CMCT in BEU buffer (50mM bicine, pH8.3, 4mM EDTA, and 7M urea) or in 30ul BEU buffer only (for mock samples) at 37°C on a thermomixer with 300rpm rotation for one hour. The CMCT reaction was stopped with 100ul of 0.3M sodium acetate and 0.1mM EDTA (pH5.6). After washing, the pellet was dried, dissolved in 40ul of 50mM sodium bicarbonate (pH10.4) at 37°C for six hours. Mock samples were handled identically in all steps but without CMCT treatment. RNA was then precipitated, washed and dissolve in RNase-free water for library preparation.

Illumina sequencing libraries were prepared following the RNA ligation method as detailed (Schwartz et al. 2014). The single-stranded cDNA product was amplified for 9-11 cycles in a PCR reaction. Libraries were pooled and sequenced on Illumina NextSeq 550 with high output of 75 bp paired end reads.

### Read trimming and mapping

Reads were adapter trimmed and quality filtered using Trimmomatic version 0.33 (Bolger et al. 2014) setting the cutoff threshold for average base quality score at 20 over a window of 3 bases, excluding the reads shorter than 20 bases post-trimming (parameters: leading=20, trailing=20, sliding window=3, Minilen= 20). Alignment of the reads to the reference sequence was done in two phases. Cleaned reads were first mapped to a sequence reference comprising of the *Drosophila melanogaster* 18S, 5.8S, 2S and 28S ribosomal RNA gene sequences (accession: M21017.1) and the sequence for RNA spike-in (Schwartz et al. 2014) using bowtie2 (Langmead and Salzberg 2012). The remaining unmapped read pairs were extracted and mapped to the *D. melanogaster* genome reference release FB2017_04 (Larkin et al. 2021) using a splice-aware aligner, STAR version 2.6.1a (Dobin et al. 2013).

### Calculation of termination ratios and termination fold change

Inserts were identified by interpolating the concordantly mapped read pairs. For each insert, the 5’ end of the read R2 was identified as the termination site of the product from reverse transcription. For each sample, a Termination Ratio (TR) was computed as the ratio of number of reads terminating at the termination site to the total coverage at the site. Also, a Termination Fold Change is computed for each site as the log2 ratio of TR in the CMC treated sample (TR_CMC_) to that in the corresponding mock treated sample (TR_Mock_).

### Identification of putative Psi sites

The termination sites with the TR_CMC_ being significantly larger than the corresponding TR_Mock_ were identified using a two-proportion z-test.

Two proportion z-test at a given nucleotide position

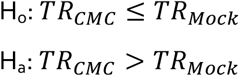

Test statistic z is calculated as

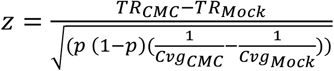

where Cvg_CMC_ and Cvg_Mock_ are the read coverage values in the CMC and mock treated samples, respectively, at the given nucleotide position and p is the pooled sample proportion which is calculated as

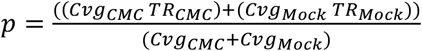

The test is considered relevant only when there are at least 5 CMC treated reads terminating at the position and another 5 continuing over the position.

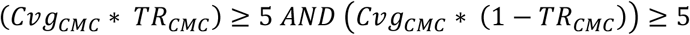

The p-values associated with the test static z (z-score) were adjusted for multiple-testing using the Benjamini and Hochberg method. Termination sites from within the annotated gene regions with Uridine as the upstream base and the adjusted p-value < 0.05 in at least two *w*^1118^ replicates, each from different experiments were considered potential Psi sites. Sites with significant adjusted p-values in 6 or 7 *w*^1118^ replicates were categorized as high reproducibility sites, those in 4 or 5 *w*^1118^ replicates as intermediate reproducibility sites and in 2 or 3 as low reproducibility sites.

### Calculation of expected number and p value of Psi-containing codons on nuclear and mitochondrial mRNAs

The probability of occurrence for a given codon with pseudouridine replacing uridine at a given position was estimated as the ratio of the number of occurrences of that codon in the coding regions of the nuclear or mitochondrial mRNA genes (with at least 10x coverage in the *w*^1118^ sample) to the total number of Uridine containing codons from those regions divided by the number of Uridines within the codon. An expected number for each Psi-associated codon was estimated by multiplying its probability of occurrence with the total number of Psi sites associated with the coding regions (907 sites associated with nuclear mRNAs and 99 sites with mitochondrial mRNAs). P-values indicate the probability of finding more than the observed number of Psi sites by chance if the observed number was greater than the expected number or the probability of finding at most the observed number of Psi sites if the observed number was less than or equal to the expected number as calculated assuming binomial distribution.

### Identification of RluA-2-dependent sites

Psi sites called using our criteria described above in *RluA-2* and *w*^*1118*^ of Experiment 2 were compared. The Psi sites that were called in at least two w^*1118*^ libraries from both of the experiments (Exp 1 and Exp 2) but not detected in any of the *RluA-2* mutant libraries are RluA-2 dependent and identified as RluA-2 targets.

### Detection of prevalent sequence motifs

For each RluA-2 dependent Psi site, a flanking sequence of 21 bp was extracted and a consensus was drawn using seqLogo version 2.8 (Crooks et al. 2004) (parameters: -B 2 -T 0.25 -C 21 -k 1 -s -10 -c -n -Y).

### Differential gene expression analysis

Mock libraries of *RluA-2* were compared against *w*^1118^ mock libraries to identify any significant differential gene expression. Read pairs mapping concordantly and uniquely to the exon regions of the annotated genes were counted using featureCounts tool ver. 2.0.0 of Subread package (Liao et al. 2014). Read alignments to antisense strand, or to multiple regions on the genome or those overlapping with multiple genes were ignored (parameters: -s 2 -p -B -C). Differential expression analysis was performed using DESeq2 ver. 1.12.3 (Love et al. 2014) and the p-values were corrected for multiple-testing using the Benjamini–Hochberg method. Genes with adjusted p-values < 0.05 were considered significantly differentially expressed.

### Gene set enrichment analysis

Gene set enrichment analysis (GSEA) was performed on a group of predefined gene sets derived from 137 KEGG (Kyoto Encyclopedia of Genes and Genomes) pathways pertaining to *Drosophila melanogaster* (Kanehisa and Goto 2000) to detect any significant, concordant differences between *RluA-2* and *w*^*1118*^ gene expression. The analysis was conducted using the software GSEA from Broad Institute (Subramanian et al. 2005) for which a matrix of DESeq2 normalized read counts for each gene were supplied as input (parameters: permute=geneset, metric= Signal2Noise, scoring_scheme=weighted).

## Accession numbers

Psi-Sequencing data have been deposited into the Gene Expression Omnibus (GEO) under the accession number GSE213312.

## Data Availability

Data generated or analyzed during this study are included in this published article (and its supplementary information files). Psi-Sequence data that support the findings of this study have been deposited in the Gene Expression Omnibus (GEO) with the accession number GSE213312.

## Acknowledgment

The authors thank the Bloomington Drosophila Stock Center (NIH P40OD018537) and Harvard Medical School for the fly stocks obtained for building the mutants used in this study. Funds to support this project were provided by the Linda and Jack Gill Center for Biomolecular Science (WDT), Indiana University startup funds (WDT), NIGMS grant 5R01GM086458 (WDT), and Indiana Clinical and Translational Sciences Institute (CTSI) 2015-2016 Postdoc Challenge award (WS).

