## Supplementary figures and images for "Transcriptome-wide analysis of pseudouridylation in *Drosophila melanogaster*"

### Figure S1

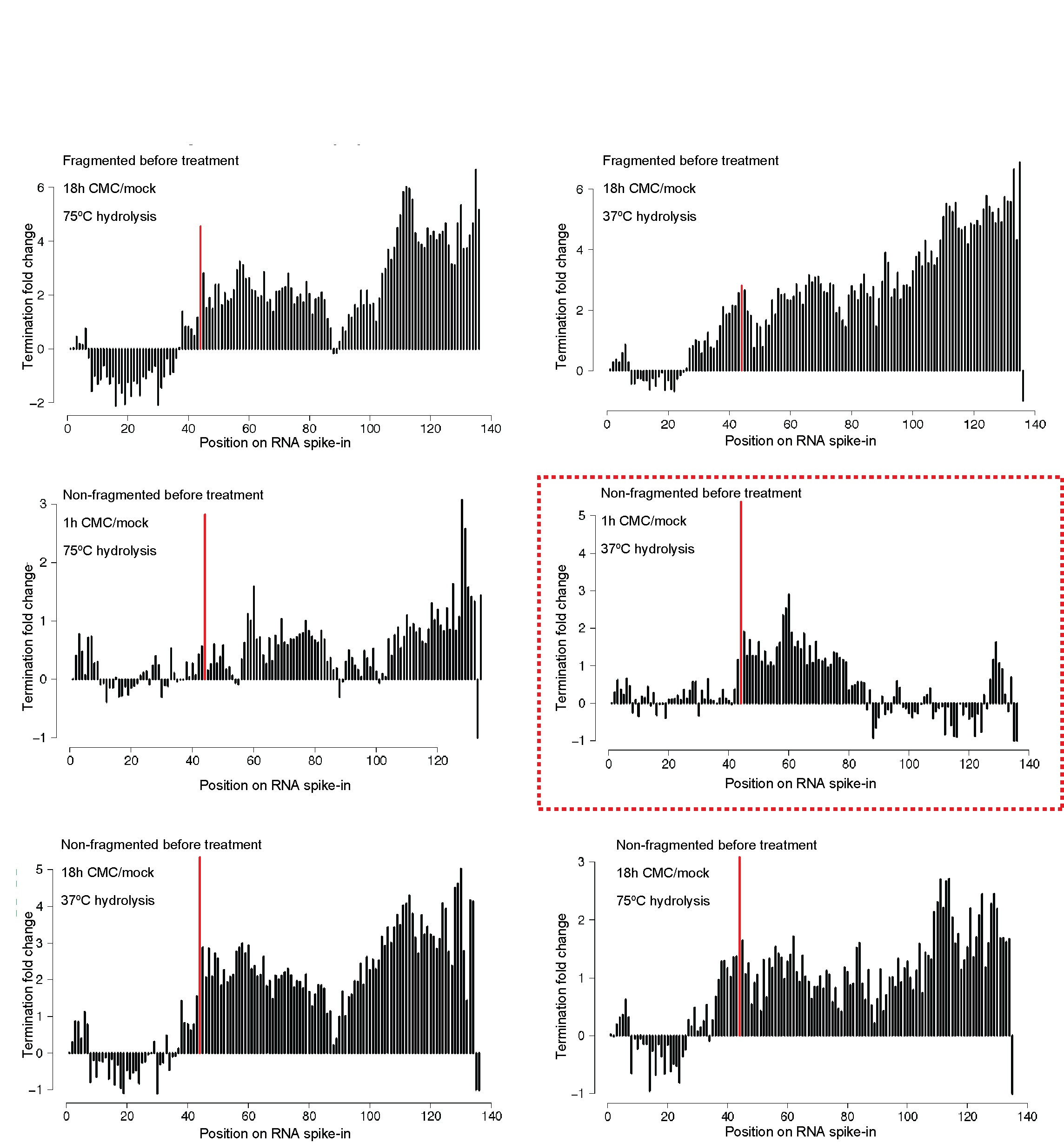

### Figure S2

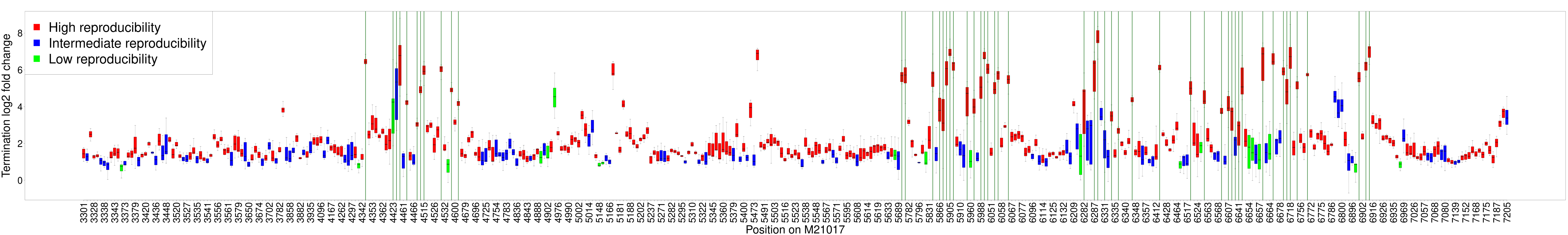

### Figure S3

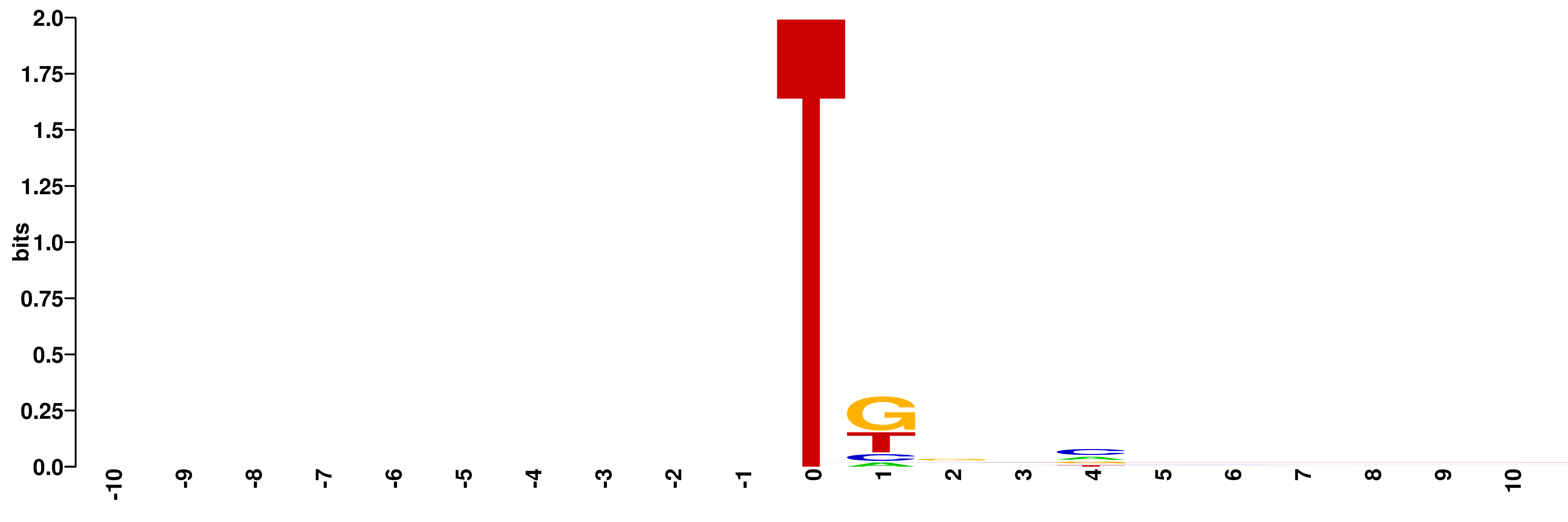

### Figure S4

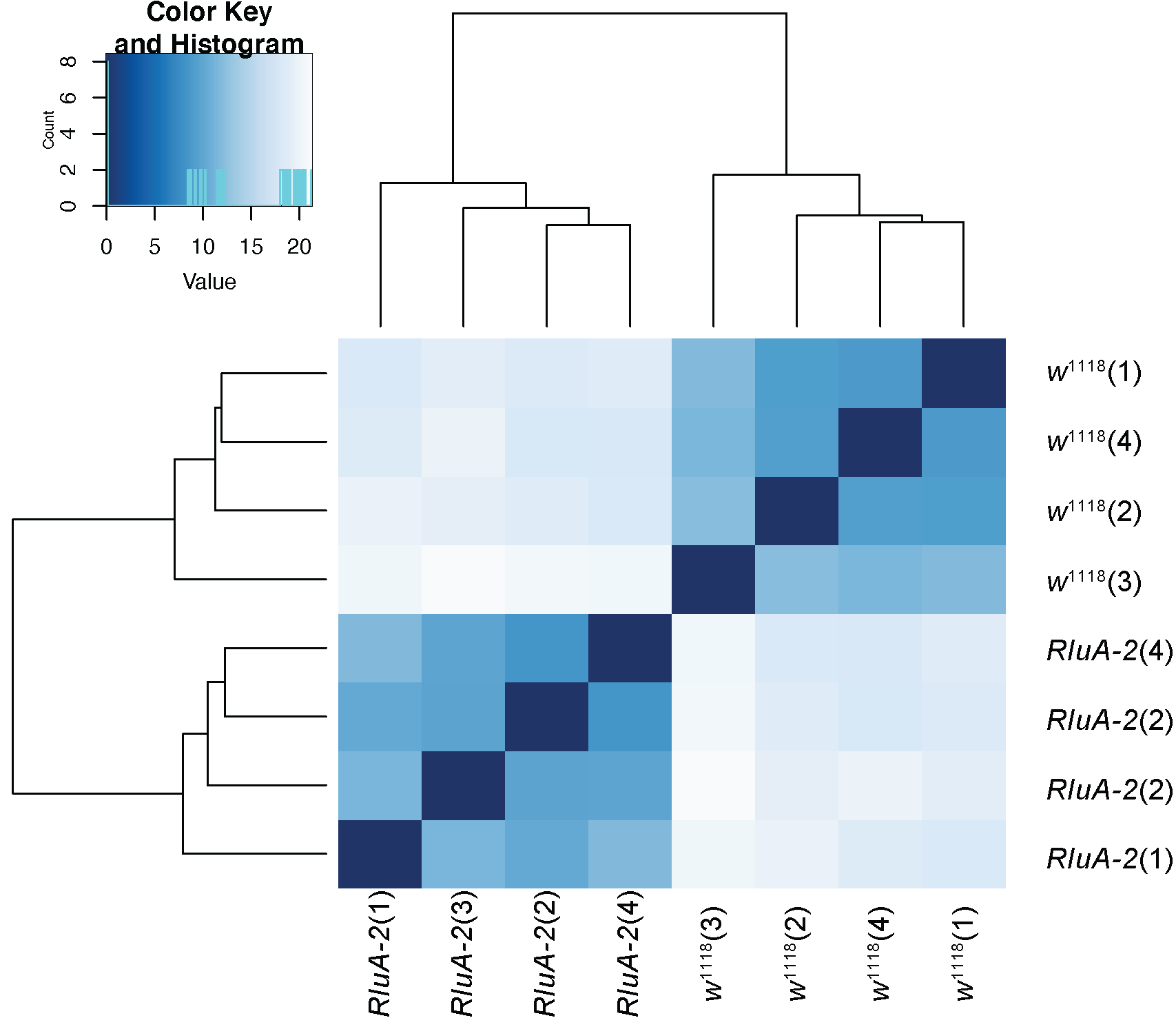
